# Distribution of contezolid in different tissues of mice and human patients infected with *Mycobacterium abscessus*

**DOI:** 10.1101/2024.12.03.626643

**Authors:** Qi Li, Yujin Wang, Qingdong Zhu, Yanrong Lin, Naihui Chu, Yu Lu, Wenjuan Nie

**Affiliations:** Department of Tuberculosis, Beijing Tuberculosis & Thoracic Tumor Research Institute, Beijing Chest Hospital affiliated to Capital Medical University, No 9,Beiguan street, Tongzhou District, Beijing 101149, P.R.China; Tuberculosis Department, Nanning No.4 People’s Hospital, No. 1 Erli, Changyi Road, Xingning District, Nanning 530012, P.R.China; Pharmaceutical Research Laboratory, Beijing Tuberculosis & Thoracic Tumor Research Institute, Beijing Chest Hospital affiliated to Capital Medical University, No 9,Beiguan street, Tongzhou District, Beijing 101149, P.R.China

**Keywords:** Contezolid, *Mycobacterium abscessus*, Drug concentration, Tissue Penetration, Pharmacokinetics

## Abstract

**Background:** Linezolid (LZD), while effective against *Mycobacterium abscessus* (MAB), can cause myelosuppression and peripheral neuropathy. Contezolid (CZD) shares a similar antimicrobial profile with improved safety, but biodistribution data remain limited. This study evaluated CZD’s biodistribution in MAB-infected mice and humans and its therapeutic potential across infection sites.

**Methods:** MICs of CZD and LZD against 32 clinical MAB isolates and three reference strains were determined. In MAB-infected mice, drug concentrations were quantified in plasma and pulmonary and cerebral tissues at 2, 4, and 8 hours post-administration. In MAB-infected patients, CZD concentrations in bone, plasma, and cerebrospinal fluid were measured at multiple time points and MAB counts in sputum cultures were assessed daily over 14 days.

**Results:** Respective MIC_50_ and MIC_90_ values were 8 and 32 µg/mL (LZD) and 16 and 32 µg/mL (CZD). Pharmacokinetic CZD and LZD level comparisons revealed peak CZD plasma levels within 2 hours, higher systemic CZD levels, comparable pulmonary tissue concentrations, and slightly lower CZD cerebral tissue penetration. In patients, CZD levels in ankle joint effusion samples reached 2.0885 µg/mL at 6 hours, peaking in the posterior malleolus. Plasma CZD concentrations peaked at 8.2349 µg/mL at 3 hours and dropped to 6.1065 µg/mL by 6 hours, while CSF levels were 0.9295 and 0.792 µg/mL at 3 and 6 hours, respectively. Sputum bacterial burden decreased rapidly within 24 hours of CZD treatment, with near-complete clearance by day 4.

**Conclusion:** CZD and LZD exhibit comparable tissue distribution but different site-specific penetration, supporting their potential for treating diverse MAB infections.

## Introduction

*Mycobacterium abscessus* (MAB) is the predominant species among rapidly growing mycobacteria (RGM), accounting for approximately 70% of RGM isolates and 25% of all non-tuberculous mycobacteria (NTM). Recognised as one of the most virulent NTM pathogens, MAB infections are associated with poor therapeutic outcomes, with treatment success rates ranging from 30% to 50%. The high prevalence and therapeutic challenges posed by MAB infections make them a significant concern in both clinical practice and research settings [1, 2].

Current treatment recommendations from the British Thoracic Society (BTS) emphasise the use of parenteral antimicrobial agents for MAB, necessitating daily visits to healthcare facilities for administration. This creates substantial barriers to treatment adherence. Additionally, prolonged treatment durations, high recurrence rates, and considerable drug-associated toxicity further complicate efforts to achieve sustained sputum culture conversion over 12 months [3-5]. Furthermore, oral therapeutic options remain limited due to restricted availability and suboptimal safety profiles.

Linezolid (LZD) is widely recommended as the preferred oral agent for MAB infections by multiple professional organisations. As an oxazolidinone antimicrobial, LZD has become a cornerstone of MAB treatment protocols. However, its clinical utility is significantly constrained by adverse events, including hematologic toxicity (e.g., myelosuppression) and neurological complications (e.g., peripheral neuropathy) [6, 7].

Contezolid (CZD) has emerged as a promising alternative, offering a comparable antibacterial spectrum to LZD but with a more favourable safety profile [8, 9]. Preliminary studies suggest that CZD exhibits potent antibacterial activity against MAB [10, 11]. Despite this potential, existing research has primarily focused on comparing the antibacterial efficacy and safety profiles of CZD and LZD, with limited exploration of their tissue distribution characteristics. Understanding tissue-specific drug exposure is essential for evaluating therapeutic efficacy, particularly for infections affecting diverse anatomical sites.

This study aimed to evaluate the *in vitro* antimicrobial activity of CZD against MAB, characterise the kinetics of its organ-specific distribution patterns in MAB-infected mice, and assess clinical outcomes of CZD treatment in selected cases of MAB infection. By integrating systemic and tissue-specific drug exposure data with *in vivo* antimicrobial activity, the findings aim to provide a foundation for the clinical implementation of CZD in managing MAB infections.

## Methods

### Minimum inhibitory concentration (MIC) determinations

The microplate alamarBlue assay (MABA) was employed to determine the LZD and CZD MICs against MAB following established standardised protocols. Antimicrobial CZD and LZD stock solutions were serially diluted to generate dilutions with concentrations ranging from 0.5 to 128 μg/mL. After the drugs were added to bacterial culture media in 96-well plates, the plates were incubated at 37 °C in a 5% CO_2_ atmosphere for 7 days. Subsequently, 70 μL of alamarBlue solution was added to each well, followed by an additional 24-h incubation. Bacterial growth was indicated by a colour change from blue to pink or purple. MIC values were defined as the lowest antimicrobial concentration that inhibited visible growth, as per Clinical and Laboratory Standards Institute (CLSI) guidelines.

### Animals

Eighteen male BALB/c mice, aged six weeks and weighing 18-20 g, were randomly assigned to two experimental groups to evaluate the pharmacokinetics of LZD and CZD. Drug concentrations were quantified at three time points (2, 4, and 8 hours post-administration), with three biological replicates per time point per group.

### Preparation of the animal model of MAB infection

Cyclophosphamide was administered intraperitoneally to induce immunosuppression at a dose of 150 mg/kg on day -4 and 100 mg/kg on day -1 before bacterial challenge. Mice were inoculated intranasally with MAB at a concentration of 4 × 10^7^ colony-forming units (CFU). CZD or LZD (50 mg/kg) was administered daily, and biological specimens were collected at predetermined time points (2, 4, and 8 hours post-administration).

### Preparation of plasma and tissue samples for drug biodistribution analysis

For tissue analysis, pulmonary and cerebral tissue specimens were mechanically homogenised. Aliquots of homogenates (10 μL) were combined with the internal standard solution, and acetonitrile was added to precipitate proteins. Samples were centrifuged, and 100 μL of the supernatant was diluted appropriately for LC-MS/MS analysis.

### Drug concentration determinations

Drug concentrations were quantified using LC-MS/MS, demonstrating linearity over the following ranges: 500– 50,000 ng/mL for plasma, 500–125,000 ng/g for pulmonary tissue, and 50–5,000 ng/g for cerebral tissue homogenates. Comparative analyses across experimental groups were performed, as presented in Tables 1 and 2.

**Table 1.** LC-MS/MS parameter settings.

|  | Q1 | Q3 | Dwell (s) | Compound | Cone (v) | Collision (eV) |
| --- | --- | --- | --- | --- | --- | --- |
| LZD | 338.1 | 296.1 | 0.108 | LZD | 25 | 17 |
| CZD | 409.1 | 269.1 | 0.108 | CZD | 55 | 28 |
| Internal standard<br>solution /Prednisone | 359.2 | 237.2 | 0.108 | Internal standard<br>solution /Prednisone | 30 | 20 |

**Table 2.** Liquid phase parameter settings.

|  |  |
| --- | --- |
| Phase A | Aqueous solution containing 0.1% formic acid |
| Phase B | Acetonitrile |
| Column temperature | 40 °C |
| Sample size | Plasma: 1 µL, tissue: 1 µL |
| Elution method | Gradient elution, 2.5 min |
| Column | Waters ACQUITY C18 1.7µm 2.1*50mm |
| Flow rate | 0.5000 mL/min |

### Determination of drug concentrations in clinical plasma samples

Plasma drug concentrations in clinical specimens were quantified using validated LC-MS/MS methodology following the same analytical protocols employed in preclinical studies. Serial plasma samples were collected at predetermined time points (2, 3, and 6 hours) following oral administration of CZD.

### Determination of drug concentrations in cerebrospinal fluid (CSF)

CZD concentrations in CSF were quantified using LC-MS/MS. Lumbar puncture was performed to obtain 0.5 mL of CSF, which was mixed with acetonitrile to precipitate proteins. The mixture was centrifuged, and the resulting supernatant was prepared for LC-MS/MS.

### Determination of drug concentrations in bone tissues

CZD concentrations in bone tissue were measured in a patient with an MAB-infected ankle joint scheduled for surgical intervention. Osseous tissue specimens (1 g) were ground and extracted with acetonitrile and methanol. The mixture was centrifuged, and the supernatant was appropriately diluted for LC-MS/MS analysis.

### Enumeration of CFU in sputa from MAB-infected patients

Two patients with MAB pulmonary infections participated in a 7-day CFU detection study. Written informed consent was obtained before study initiation, and participants were hospitalised throughout the study period.

At enrollment, baseline clinical and microbiological assessments were conducted. Pharmacokinetic/pharmacodynamic (PK/PD) sampling involved serial blood draws at 2 hours post-dose on days 7 and 21 of therapy. Expectorated sputum specimens were collected following standardised protocols:

1. Overnight sputum samples were obtained following mouth rinsing with sterile saline.
2. Samples were stored at 4–8 °C under controlled conditions.
3. Temperature-monitored transport protocols ensured sample integrity.

### CFU enumeration

Quantitative microbiological analysis was performed on sputum specimens using serial dilution techniques. Aliquots were inoculated onto Middlebrook 7H10 agar medium and incubated aerobically at 37 °C for 7–14 days. CFU were enumerated following established protocols, with bacterial counts were calculated based on dilution factors and expressed as CFU/mL.

## Results

MIC values of LZD and CZD were evaluated against 32 clinical MAB isolates and 3 reference strains (Table 3). For LZD, the average MIC_50_ and MIC_90_ values of the isolates were 8 µg/mL and 32 µg/mL, respectively, with MIC values ranging from 4 to 64 µg/mL. In comparison, CZD exhibited MIC_50_ and MIC_90_ values of 16 µg/mL and 32 µg/mL, respectively, with a distribution range of 2-64 µg/mL.

**Table 3.** MIC value of LZD and CZD against MAB.

|  | MIC<sub>50</sub>(ug/mL) | MIC<sub>90</sub>(ug/mL) | MIC range(ug/mL) |
| --- | --- | --- | --- |
| LZD | 8 | 32 | 4-64 |
| CZD | 16 | 32 | 2-64 |

Following oral administration in mice, CZD and LZD exhibited peak concentrations in plasma, pulmonary tissue, and CSF at 2 hours post-dose, followed by a time-dependent decline (Figure 1, Table 4). A comparative pharmacokinetics analysis revealed distinct tissue-specific distribution patterns. In plasma, CZD demonstrated significantly higher concentrations than LZD at 2 hours post-administration (*P* < 0.05), with this trend persisting but not reaching statistical significance at 4 hours (*P* = 0.3921). By 8 hours, the CZD concentration was marginally lower than that of LZD (*P* = 0.6609).

**Table 4.** Temporal distribution of CZD in multiple tissue compartments of MAB-infected mice.

| | CZD<br>(mean ± variance) | LZD<br>(mean ± variance) | $P$ |
| --- | --- | --- | --- |
| Plasma (ug/mL) |  |  |  |
| 2 h | 24.9 ± 7.28 | 18.2 ± 1.2067 | < 0.05 |
| 4 h | 11.30 ± 0.65 | 10.45 ± 0.914 | 0.3921 |
| 8 h | 5.07 ± 0.94 | 5.85 ± 5.853 | 0.6609 |
| Pulmonary tissue<br>(ug/mL) |  |  |  |
| 2 h | 20.10 ± 1.93 | 21.03 ± 0.549 | 0.4487 |
| 4 h | 10.95 ± 8.82 | 11.37 ± 0.336 | 0.8562 |
| 8 h | 3.76 ± 0.49 | 5.68 ± 5.4434 | 0.3296 |
| Cerebrospinal fluid<br>(ug/mL) |  |  |  |
| 2 h | 2.53 ± 0.0953 | 3.93 ± 0.0622 | 0.0075 |
| 4 h | 1.19 ± 0.108 | 2.08 ± 0.0388 | 0.0296 |
| 8 h | 0.35 ± 0.0015 | 1.39 ± 0.0025 | 0.0003 |

**Figure 1.**
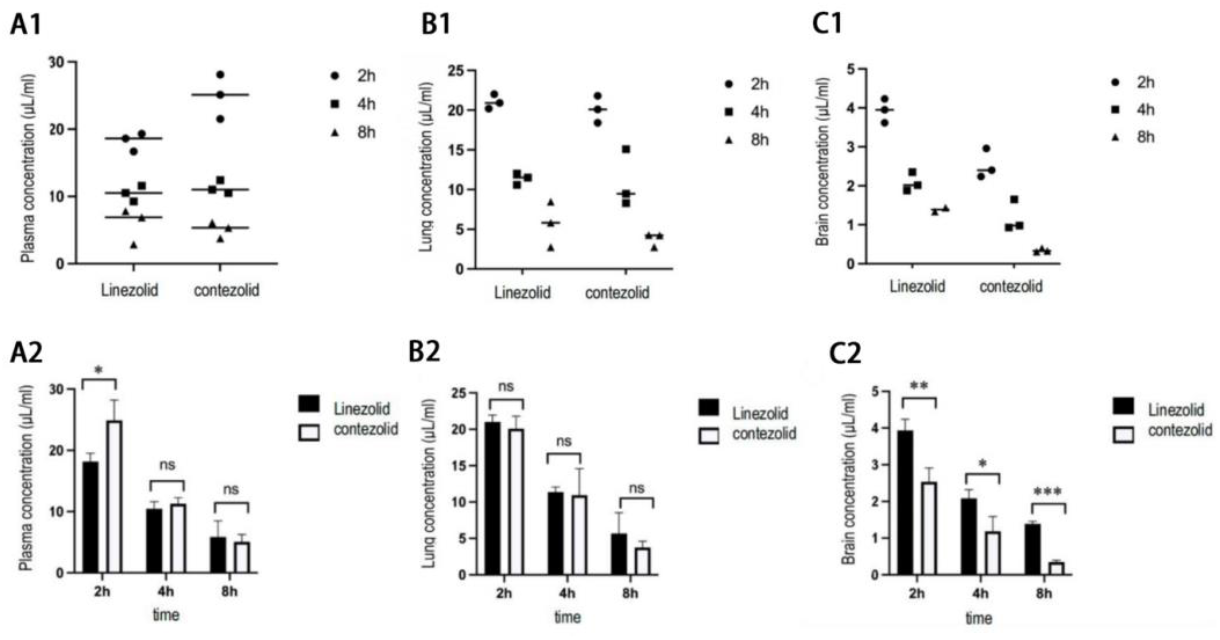
Drug concentrations at different sites in a mouse model of MAB infection. A: Plasma drug concentration; B: Pulmonary tissue drug concentration; C: Cerebrospinal fluid drug concentration. Each group contains three mice. \**P* < 0.05, \*\**P* < 0.01.

In pulmonary tissue (Figure 1, B1&2), LZD exhibited slightly higher concentrations across all sampling time points compared to CZD, though the observed differences did not achieve statistical significance (2 hours: *P* = 0.4487; 4 hours: *P* = 0.8562; 8 hours: *P* = 0.3296).

The analysis of CSF drug concentrations (Figure 1, C1&2) revealed significantly elevated LZD levels compared to CZD at all time points (2 hours: *P* = 0.0075; 4 hours: *P* = 0.0296; 8 hours: *P* = 0.0003). Notably, both compounds demonstrated significantly higher plasma and pulmonary tissue concentrations compared to CSF.

The patient underwent surgical intervention on the ankle joint 6 hours after oral CZD administration. The concentration of CZD in the ankle joint effusion was determined to be 2.0885 µg/mL. Differential analysis of osseous tissue revealed a concentration gradient across anatomical sites, with the highest levels observed in the posterior malleolus, followed by the medial and lateral malleoli, and the lowest concentrations in articular surface tissues. This spatial distribution pattern may correlate with the regional vascularity of the ankle joint complex (Table 5).

**Table 5:** Site-specific concentrations of CZD in ankle joint components at 6 hours post-oral administration.

| Site of the ankle joint | CZD concentration(ug/mL) |
| --- | --- |
| Ankle joint effusion | 2.0885 |
| Articular surface | 0.0000 |
| medial malleolus | 0.2697 |
| lateral malleolus | 0.1779 |
| posterior malleolus | 0.3384 |
| quality control (QC) | 10.2070 |

Pharmacokinetic analysis of clinical specimens (Table 6) demonstrated that plasma CZD concentrations reached 8.2349 μg/mL at 3 hours post-administration, subsequently declining to 6.1065 μg/mL at 6 hours. CZD concentrations in CSF were 0.9295 μg/mL and 0.792 μg/mL at 3 and 6 hours, respectively. The CSF-to-plasma concentration ratios remained relatively stable over time, indicating consistent but limited blood-brain barrier (BBB) penetration. These findings align with penetration characteristics observed in preclinical mouse models, confirming CZD’s moderate central nervous system bioavailability.

**Table 6:** Temporal profile of CZD concentrations in plasma and CSF of an MAB-infected patient.

| Plasma |  | CSF |  |
| --- | --- | --- | --- |
| 3h | 8.2349 ug/mL | 3h | 0.9295ug/mL |
| 6h | 6.1065 ug/mL | 6h | 0.792ug/mL |

A quantitative microbiological analysis of serial sputum specimens was performed in two patients with confirmed MAB pulmonary infection undergoing CZD therapy. A significant reduction in bacterial burden, as indicated by CFU counts, was observed within 24 hours of treatment initiation. By day 4 of oral CZD administration, sputum specimens exhibited near-complete bacterial clearance (Table 7).

**Table 7:** The quantitative analysis of MAB burden in serial sputum specimens: Log10 CFU measurements.

| No.\Times | D1 | D2 | D3 | D4 | D7 |
| --- | --- | --- | --- | --- | --- |
| No.1 | 2.605 | 2.358 | 1.65 | 0.603 | 0 |
| No.2 | 1.17 | 0.845 | 0 | 0 | 0 |

## Discussion

The prototype oxazolidinone antibiotic LZD is effective against MAB infections across various anatomical sites due to its favourable tissue distribution profile and ability to penetrate the BBB. However, LZD’s molecular target, the bacterial ribosome, shares structural homology with mitochondrial ribosomes in human somatic cells, leading to significant adverse reactions such as myelosuppression and peripheral neuropathy. Prolonged LZD therapy, particularly in combination regimens, increases the likelihood of toxic effects, resulting in treatment cessation and poor therapeutic outcomes.

Contezolid (CZD), structurally distinct from LZD due to the incorporation of three fluorine atom substituents into its B ring, exhibits a non-coplanar structure that may enhance its safety profile. A clinical study reported that 90% of patients experiencing LZD-associated adverse events showed improvement or resolution of symptoms within one month of transitioning to CZD-containing regimens [9]. Preclinical studies have shown that both LZD and CZD exhibit comparable inhibitory effects on intracellular mycobacterial proliferation [10].

Given CZD’s improved safety profile compared to LZD, this study aimed to evaluate its antimicrobial activity against MAB both *in vitro* and *in vivo*, as well as its tissue distribution kinetics, to provide foundational pharmacokinetic data for optimising therapeutic strategies. MIC testing of 32 clinical isolates and three reference strains revealed comparable MIC distributions between CZD and LZD, with CZD displaying a broader range of activity. These findings align with previous studies documenting similar *in vitro* antimicrobial efficacy between the two drugs across 194 clinical isolates [11].

The established therapeutic efficacy of LZD is attributed to its favourable tissue distribution. Although CZD displays comparable *in vitro* antimicrobial activity to LZD, evaluating its *in vivo* tissue distribution profile is critical to understanding its target site accessibility and therapeutic potential. CZD demonstrated optimal plasma exposure, achieving peak concentrations in synchrony with LZD and displaying comparable concentration-time profiles. In pulmonary tissue, CZD and LZD concentrations were similar, reflecting the extensive vascularisation of the lungs.

However, both compounds exhibited significantly lower concentrations in cerebral tissue compared to plasma and pulmonary compartments, likely due to BBB constraints. CZD’s limited CNS penetration may be attributed, at least in part, to its high-level binding to plasma proteins, which restricts the unbound fraction available to cross the BBB.

Although CZD exhibited lower pulmonary concentrations at all time points, these differences were not statistically significant. Conversely, CZD’s penetration into CSF was significantly lower than LZD (P < 0.05), indicating limited CNS bioavailability. Despite this, CZD achieved concentrations in plasma, pulmonary tissue, and bone tissue that exceeded the MIC for MAB at 2 hours post-administration, supporting its efficacy in highly perfused tissues. However, clinical CSF concentrations remained below the MIC, suggesting that CZD alone may be insufficient for clearing CNS infections.

Clinical pharmacokinetic data largely corroborated preclinical findings, with plasma concentrations significantly higher than those in CSF. Plasma levels peaked at 2 hours post-administration while CSF concentrations peaked at 3 hours, aligning with a previous study showing that clinical CSF concentrations of CZD remained below the MIC even at peak levels (3 hours) relative to the established MIC range (0.5–64 µg/mL) [11].

Osseous tissue drug concentrations, particularly in the ankle joint, were lower than in plasma and CSF, potentially reflecting differences in regional vascularity. Nonetheless, CZD concentrations in specific bone regions exceeded the MIC, suggesting its potential for treating bone infections. Preoperative fasting protocols likely impaired drug absorption, contributing to suboptimal osseous tissue concentrations. Enhanced CZD bioavailability under high-fat dietary conditions has been documented in previous studies, suggesting that therapeutic bone tissue levels may be achievable under normal dietary conditions. Additionally, infection-associated inflammation may impair regional perfusion, further limiting drug penetration into affected osseous tissues.

The inability to assess pulmonary tissue distribution in clinical subjects due to ethical and procedural constraints related to repeated bronchoalveolar lavage fluid collection is a limitation of this study. However, preclinical data suggest superior pulmonary versus cerebral tissue penetration for CZD, consistent with the greater vascularisation of lung tissue. These findings suggest that therapeutic drug concentrations may be achievable in human pulmonary tissue.

While clinical evidence for CZD’s efficacy against MAB infections remains limited, a case report described the successful treatment of a skin MAB infection in a 38-year-old female patient. Following a 6-month course of CZD-containing therapy, the patient’s lesions gradually shrunk and fully resolved without recurrence by 10 months post-treatment [12]. In the current study, CZD demonstrated early bactericidal activity against pulmonary MAB infections, with significant reductions in sputum bacterial burden and near-complete clearance by day 4 of treatment.

To expand the clinical evidence base, this investigation evaluated the early bactericidal activity of CZD monotherapy in pulmonary MAB infection through quantitative sputum culture analysis. Despite the limited cohort size, the data demonstrated promising antimicrobial activity, as evidenced by a significant reduction in sputum bacterial burden.

These findings suggest that CZD may serve as a promising alternative to LZD for treating pulmonary, skin, and soft tissue MAB infections. However, for CNS infections, combination therapy with other antimycobacterial agents may be necessary to achieve adequate therapeutic drug exposure at target sites. Limitations of this study include the small clinical sample size, restricting the generalisability of tissue distribution findings. Larger clinical studies are needed to establish robust pharmacokinetic profiles and optimise therapeutic applications. Additionally, ethical constraints precluded an assessment of CZD’s pulmonary tissue disposition in clinical subjects, a critical area for future research given that pulmonary infections are the primary manifestation of MAB. Lastly, this study did not evaluate organ-specific pathological changes, *in vivo* antimicrobial activity, or metabolite-associated toxicity in preclinical settings, highlighting opportunities for future investigation.

In conclusion, CZD exhibits comparable antimicrobial activity to LZD with a more favourable safety profile, supporting its potential as a therapeutic alternative for MAB infections. However, further studies are warranted to address its limitations and fully elucidate its clinical efficacy.

